# A deep learning and digital archaeology approach for mosquito repellent discovery

**DOI:** 10.1101/2022.09.01.504601

**Authors:** Jennifer N. Wei, Carlos Ruiz, Marnix Vlot, Benjamin Sanchez-Lengeling, Brian K. Lee, Luuk Berning, Martijn W. Vos, Rob W.M. Henderson, Wesley W. Qian, D. Michael Ando, Kurt M. Groetsch, Richard C. Gerkin, Alexander B. Wiltschko, Jeffrey Riffel, Koen J. Dechering

## Abstract

Insect-borne diseases kill >0.5 million people annually. Currently available repellents for personal or household protection are limited in their efficacy, applicability, and safety profile. Here, we describe a machine-learning-driven high-throughput method for the discovery of novel repellent molecules. To achieve this, we digitized a large, historic dataset containing ∼19,000 mosquito repellency measurements. We then trained a graph neural network (GNN) to map molecular structure and repellency. We applied this model to select 317 candidate molecules to test in parallelizable behavioral assays, quantifying repellency in multiple pest species and in follow-up trials with human volunteers. The GNN approach outperformed a chemoinformatic model and produced a hit rate that increased with training data size, suggesting that both model innovation and novel data collection were integral to predictive accuracy. We identified >10 molecules with repellency similar to or greater than the most widely used repellents. We analyzed the neural responses from the mosquito antennal (olfactory) lobe to selected repellents and found a limited correlation between these responses and our GNN representation. This approach enables computational screening of billions of possible molecules to identify empirically tractable numbers of candidate repellents, leading to accelerated progress towards solving a global health challenge.

## Introduction

Mosquitos and other blood-sucking arthropods carry and transmit diseases that kill hundreds of thousands of people each year^1,2^. To make continued progress on this global health issue, we must discover, manufacture, and deploy more efficient molecules for pest control across a variety of application spaces collectively termed “vector control”; this includes molecules that affect life history traits, such as insecticides, and molecules that affect host-seeking behavior, e.g. topical repellents for personal protection and spatial repellents applied to a home or room. Commonly used repellents such as DEET (N,N-diethyl-meta-toluamide), Picaridin (Hydroxyethyl isobutyl piperidine carboxylate), and IR3535 (Ethyl butylacetylaminopropionate) require high concentrations of over 40% \cite{noauthor_undated-nv} which limit their use to topical applications. Furthermore, they have undesirable properties and/or safety profiles; for example, DEET is a plasticizer, precluding its use on synthetic clothing or shelter surfaces, and it is toxic to some vertebrate wildlife^3^. Some commonly used repellents are species-specific; for example IR3535 is effective against *Aedes aegypti* but is ineffective against *Anopheles* mosquitoes and is therefore not recommended for use in malaria-endemic regions. Over the past few decades, only a few dozen new repellent molecule candidates have been found and very few have reached the market; an approach to rapidly discover and validate large numbers of new candidates is desperately needed.

Multiple strategies exist for identifying insect repellent candidates. Behavioral assays seek to directly test repellent activity in realistic conditions. Recognizing the devastating effect of insect-borne diseases (including dengue fever) faced by the United States Army during the second world war, the U.S. Department of Agriculture (USDA) tested 30,000 molecules for their effectiveness as repellents and insects on mosquitos, ticks, and other insect species^4,5^. In particular, 14,000 molecules were tested for their effectiveness as mosquito (*A. aegypti* and *Anopheles quadrimaculatus*) repellents using human volunteers; this effort led to the discovery of DEET. Structure-targeted modeling of the obligatory insect olfactory co-receptor Orco led to the discovery of VUAA1^7^. Scaffold-hopping techniques^8^ can focus the molecular search space, and in combination with arm-in-cage testing, led to the discovery of IR3535^9^ and DEPA^10^. Chemoreceptor studies exploit the molecular mechanism of action: DEET and IR3535 modulate the activity of odorant and gustatory receptors^11,12^ but may also affect cholinergic signaling^13,14^. The exact molecular details of their mode of action are not fully understood, and may be very species-specific (Afify and Potter, 2020). It is difficult to more broadly and systematically explore molecular space using each of these approaches, as they can be labor-intensive.

The USDA dataset represents a wealth of information on the relationship between molecular structure and arthropod behavior. Small parts of this dataset have been used previously to train computational models of mosquito repellency^15–17^, typically on specific structural families of molecules. Katritzky et al.^18^ used an artificial neural network model trained on 167 carboxamides and found 1 carboxamide candidate with high repellency activity. As modern deep learning models show performance which scales in proportion to the volume of their training data^19^, we hypothesized that exploiting the full size of the USDA dataset would provide a strong starting point for a new deep learning model. We selected a graph neural network architecture (GNN), as GNNs have been shown to have superior performance to computable cheminformatics descriptors in predicting the properties of a molecule from its chemical structure, given a sufficiently large dataset^20,21^. Notably, previous work demonstrated that a GNN-based human odor model outperforms standard cheminformatics models even on insect behavior datasets.^15–17^

Here we present a data-driven workflow for the discovery and validation of novel molecules for behavioral modification in arthropods. The critical components underlying the success of this approach are 1) expanded training data made possible by a complete digitization of the USDA dataset; 2) high-quality validation data using a parallelizable membrane-feeding assay that does not require human volunteers; and 3) a graph neural network model to learn the relationship between molecular structure and these data. We iteratively use this model to propose candidates from a purchasable chemical library, validate these candidates for repellency, and use these results to expand the training dataset and therefore improve the predictive accuracy of the behavior model (Figure 1). We performed calcium imaging experiments to measure mosquito neural activity, to search for patterns in the mosquito’s perception of our selected compounds and our graph neural network representations. Through this process, we have discovered a chemically diverse set of molecules with effectiveness equal to or greater than DEET, unlocking new potential capabilities in vector control.

**Figure 1:**
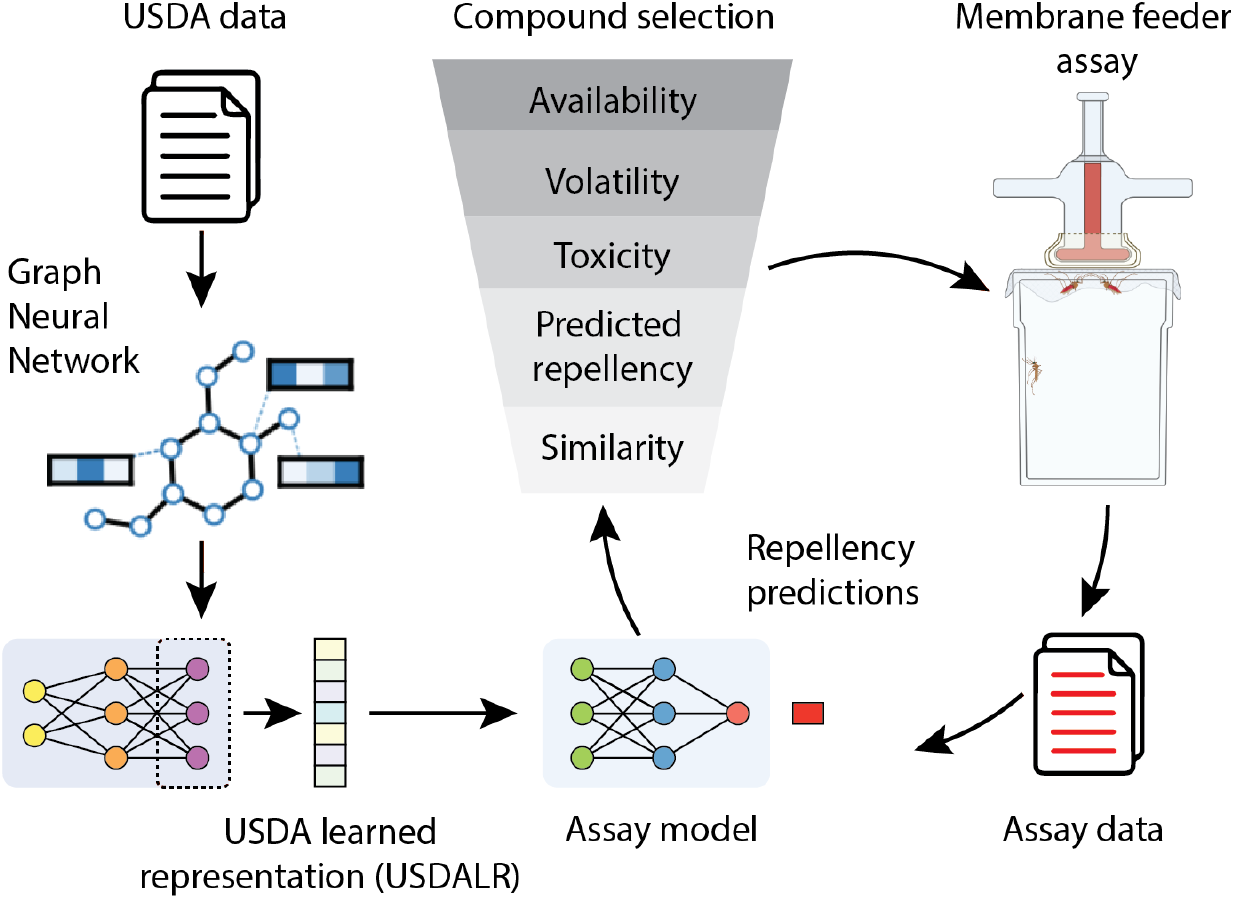
Pipeline for active learning of new behavior repellent molecules. A large historical dataset from the USDA (*USDA data*) was used to train a graph neural network to generate a fixed vector representation of any candidate molecule (*USDA learned representation, USDALR*). To create the transfer-learned *assay model*, molecules are first embedded with the USDA learned representation and fed to a dense neural network; this assay model is trained on the *assay data*. A large-scale *in silico* molecular screen is applied to select candidate molecules for testing in a membrane feeder assay for repellency. Resulting data are used to train the assay model. In subsequent iterations, the assay results are used to improve the transfer-learning model, a form of active learning.

## Results

### Digitizing a rich historical dataset

The USDA dataset is unmatched in size and scope, but for decades existed only in print. Google Books scanned and made available the original work online^4^, and for this work we subsequently converted it into a machine-readable format. After some preprocessing to make the dataset easier to read, we employed expert curators to transcribe the full records and provide canonical structures for each listed molecule (Fig. 2A, Methods). We then focused our analysis on the four mosquito repellency assays contained in this dataset: two mosquito species, *A. aegypti* and *A. quadrimaculatus*; and two repellency contexts, skin and cloth. Together these comprise ∼19,000 labeled data points on repellency of specific molecules (Fig. 2B), representing a broad range of structural and functional classes (Fig. 2C). This large dataset served as training data for our modeling efforts.

**Figure 2:**
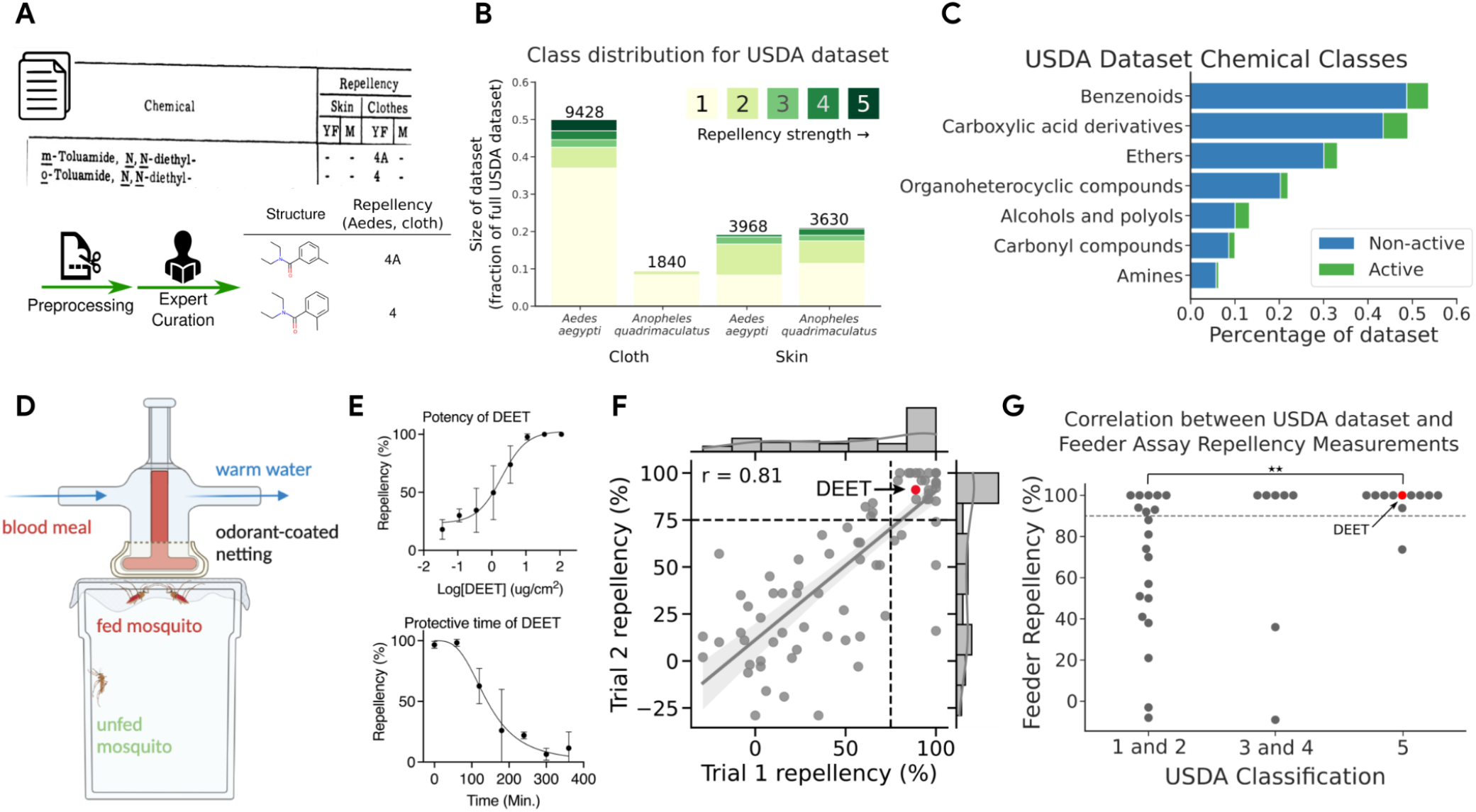
Overview of data sources. **(A)** The USDA dataset scanned into Google Books was digitized and manually curated into a machine-readable table of repellency ratings for each compound (King, WV 1954). **(B)** Digitized ratings from USDA dataset used here covered two assay types and two mosquito species. **(C)** The USDA dataset covered a diverse range of chemical classes; shown here is the distribution of some ClassyFire classes (Djoumbou-Feunang et al. 2016). Active compounds are defined as class 4 or higher. **(D)** Our validation assay used warmed blood and an odorant-coated netting; repellency was identified with a decrease in feeding behavior relative to a control odorant (ethanol). **(E)** Repellency measured using the assay in (D); 100% indicates total repellency (no feeding) and 0% matches behavior using the solvent alone. Data points (mean +/-SD across replicates) show repellency using the indicated concentration of DEET as the odorant. Top: Repellency of DEET at t=120 min. increases with concentration. Bottom: Repellency decreases with time after initial application of the odorant (sigmoidal fit). **(F)** Repellency values are correlated across independent replications of the assay. Trials 1 and 2 are not necessarily in chronological order. Test-retest values of DEET are indicated in red. Dotted line indicates positive activity cutoff at Repellency=0.75 for t=120min. (**G**) Repellency observed in the assay at t=2 min. at 1% concentration using *A. stephensi* is concordant with repellency from the USDA dataset using *A. aegypti* on cloth. Dotted line represents activity cutoff at Repellency=0.9 for t=2min. for feeder assay. DEET’s activity is represented by a red dot. Raw repellency % for USDA Class 1&2 vs Class 5: p<0.01 (Mann-Whitney U Test); Hit percentage: p<0.05 (Z-test of proportions).

### Assessment of repellent candidates

In order to test model predictions and iteratively expand the training data, we adapted a standard membrane feeding assay (SMFA), commonly used in malaria research^22,23^, to evaluate the repellency against *Anopheles stephensi* mosquitoes. Repellency was evaluated by prevention of blood feeding relative to a vehicle (ethanol) control (Fig. 2D). The assay was used to evaluate each molecule’s potency and duration of effect as exemplified for the reference molecule DEET in Fig 2E. We assessed the inherent inter-assay reliability by comparing repellency levels for a diverse set of molecules from independent experiments (tested at 25 µg/cm^2^, r=0.81, Fig. 2F). Using a cut-off of 75% repellency as measured 120 min after initial application, selected to include widely used repellents (e.g. DEET, dimethyl phthalate, and indalone), approximately 3/4 of the molecules classified as active in a first assay were confirmed to be active upon re-testing.

The USDA dataset was collected ∼70 years ago using arm-in-cage experiments, involving human volunteers, while our assay was conducted with a surrogate target. We evaluated the relationship between these two experiments by directly comparing the activity of 38 molecules with their repellency reported in the USDA dataset. We found considerable concordance between the historical USDA dataset and the membrane feeding assays (p<0.01 Mann Whitney U test, Fig. 2G), despite differences in experimental setup. However, some disagreement was observed, highlighting the need for additional data collection.

### Modeling mosquito repellency behavior

Using the USDA dataset, we sought to create a representation of molecules specific to mosquito repellency behavior. It has been previously demonstrated that graph neural networks (GNNs) are particularly adept at creating task-specific representations^20,24^, and that representational power extends to the domain of olfaction^25,26^. We trained GNN models on the USDA dataset, observing an AUC=0.881 on the cloth-*Aedes aegypt*i task, the task with the largest dataset (Methods). We then use the output heads from the ensemble models on all four USDA tasks to create the *USDA learned representation* (USDALR, Figure 1).

We sought to build a model that was specific for the activity behavior in our membrane feeder assay. We created an *assay model* by first using the fixed USDA learned representation to embed input molecules, then adding a two layer, 256-node neural network to learn to predict the assay data.

We applied the assay model to make predictions on novel repellent candidate compounds from a large library of purchasable molecules provided by the vendor eMolecules^27^. We filtered this library for desirable qualities such as volatility and low cost, and we further screened out molecules which did not pass an inhalation toxicity filter (Methods). From among those compounds passing these filters (∼10k molecules), we selected those which had sufficient predicted repellency and--to ensure novelty--which were structurally distinct (Tanimoto similarity <0.8) from those in the USDA dataset or previous candidate selections. Assay results from each batch of selections were added to the assay dataset; for each subsequent batch of selections, the assay model was re-trained on the expanded assay dataset. Detailed notes on the specific modeling setup for each batch are located in the Supplementary section.

Over several iterations, a total of 400 molecules were purchased and further screened empirically according to a solubility criterion (Methods); those that passed (n=317) were then tested for repellency with the membrane-feeder assay. Over the course of selections spanning over a year, some adjustments were made to both the USDA model and the membrane-feeder assay. In particular, our hit definition evolved with our dataset size and model capability: we initially defined a hit as ≥90% repellency using a dose of 25 µg/cm^2^ as measured at T=2min (≥1 measurement), but in the final batch of selections, we changed our definition to ≥75% repellency as measured at T=120min (≥3 measurements).

### The hit rate improves with training data size

To evaluate the contribution of the training data to our performance, we retrospectively scored high-repellency candidates in two phases: before the USDA dataset was available (pre-USDA) and after we began using the USDA dataset to build and deploy the USDA learned representation (post-USDA). In the pre-USDA phase, instead of using the USDA learned representation to embed molecules, we employed an odor-specific representation previously demonstrated to outperform standard cheminformatics representations on olfaction-related tasks^26^. Further, at that time, we only had assay data for 34 molecules, so we opted to use a k-nearest neighbors model (k=10) to model assay activity. In the post-USDA phase, the assay dataset size for the first batch was 142 molecules, and grew to a size of 402 molecules for our final batch of selections (Supplemental Batch Notes).

This large dataset made a huge difference; hit rates post-USDA measured on repellency time=2min increased to 49% from the pre-USDA level of only 29% (Figure 3A). When we then raised the bar for “hit” classification to require a longer duration of effect, hit rates dropped to 6% for predictions from the post-UDSA phase and 3% for predictions from the pre-USDA phase. It is important to note that only the *last* batch in the post-USDA phase was trained to find candidates meeting this new repellent standard; further iterations may have continued to improve performance as they did under the previous standard.

**Figure 3:**
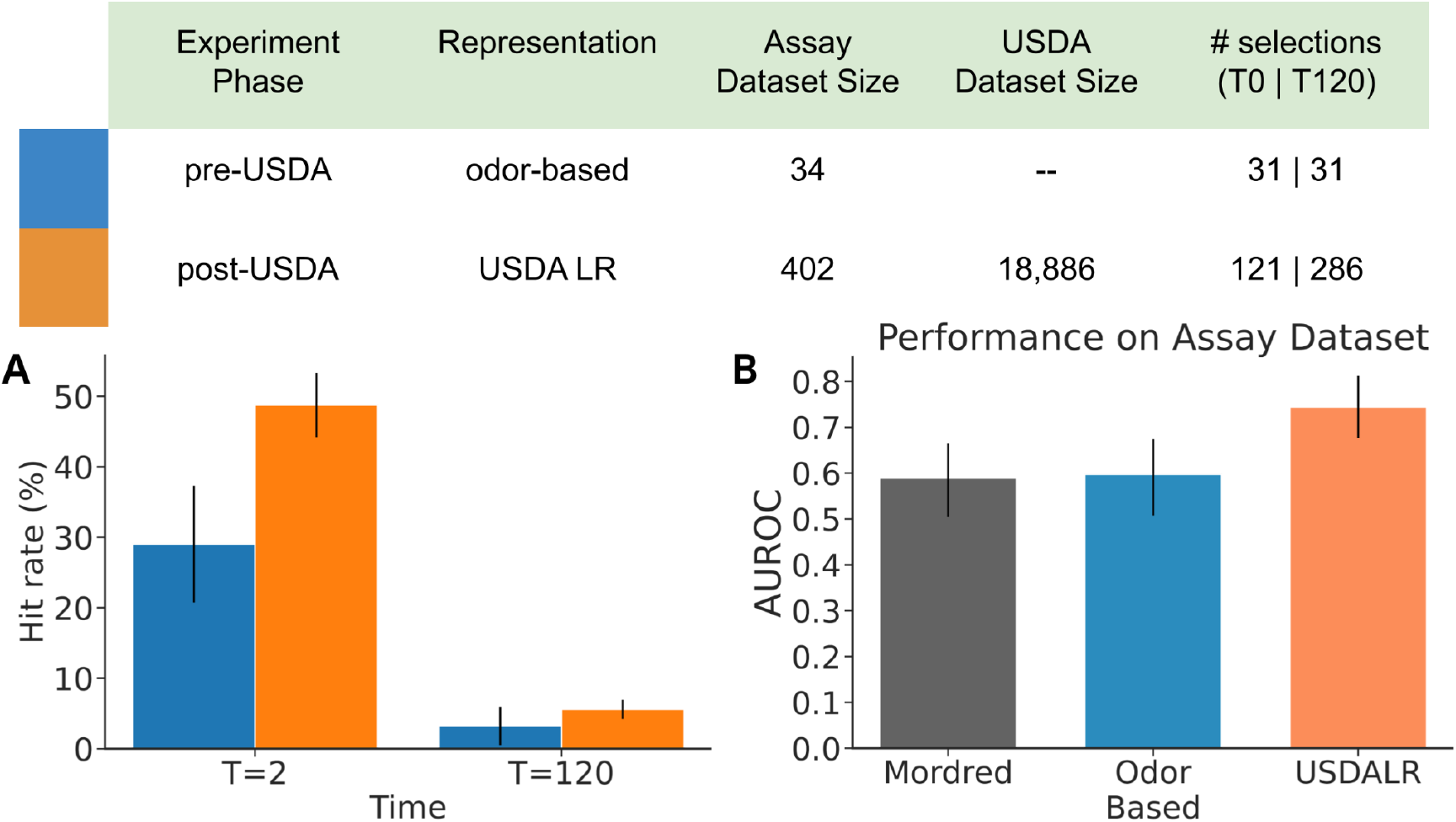
The table reflects experimental testing set up in pre-USDA phase, i.e. before the use of the USDA dataset for modeling, and post-USDA phase, i.e. after the use of the USDA dataset. **(A)** Active repellent compounds found at a much higher rate in post-USDA phase (49%) vs. pre-USDA phase (29%). Hits are defined as compounds that showed >90% repellency in the feeder assay at initial application (t=2 min) or >75% repellency after 2 hours of evaporation (t=120). Error bars represent the standard error of jackknife estimated mean values. **(B)** In a retrospective prediction task, USDA learned representation model (USDALR) outperforms models using cheminformatics representation (Mordred, Moriwaki et al, 2018) and odor-based representation (Qian et al. 2022). Models were trained on assay data collected before USDA modeling (88 data points), and evaluated on post-USDA measurements (170 data points). Error bars represent 95% bootstrap-resampling confidence intervals.

This “hit rate” comparison across the two different experimental phases aggregates changes in both representational approach and assay dataset size; how much did the USDA learned representation specifically, and by extension the USDA dataset, improve our model’s performance?

To estimate the contributions of the USDA representation, we performed a retrospective analysis comparing the USDA representation against two other chemical representation approaches: a cheminformatics representation (using Mordred descriptors^28^) and the odor-based representation^26^ used in the pre-USDA phase. The same assay model architecture was used for the different representations. We split the full assay dataset into two parts, a training set composed of molecules from all batches of tests performed before the use of the USDA dataset (88 measurements) and an evaluation set of all molecules selected in the post-USDA phase (170 measurements).

We observed that the USDA learned representation model significantly outperformed both alternatives on this prediction task (Figure 3B; USDA model AUC=0.74 [0.68,0.81]; Chemoinformatics model AUC=0.59 [0.50,0.67]; GNN Odor model AUC=0.60 [0.51,0.67]), suggesting that the historical dataset played a significant role in the elevated predictive performance. There is a selection bias because the selection of molecules for evaluation was done by the assay model using USDA learned representations. One effect of this bias is that it reduces the expected number of negative examples, reducing the contrast between predicted repellents and non-repellents, resulting in a *negative* bias into all AUC measurements. However, the model used for selection should suffer the greatest negative bias, suggesting that the performance difference we observed is an underestimate of the true advantage that the USDA model has over its alternatives, as would have been observed under a counterfactual unbiased selection of repellent candidates.

### Selected hit molecules are chemically diverse

Training a model on a large pool of data containing a variety of molecules allows the model to generalize to larger areas of chemical space. Figure 4 shows the distribution of molecules selected by our post-USDA models, and compares them to the active molecules reported in the USDA dataset itself. The candidate selections made by our model explore some of the same regions of the USDA dataset, but find hits in some underexplored regions of the original dataset (Figure 4A). The ML-selected molecules were required to be a minimum of 0.2 Tanimoto distance from USDA molecules; we observe an overall median Tanimoto distance of 0.52 from USDA molecules across all of our selections, and a median distance of 0.48 from USDA molecules amongst active molecules (Figure 4B). Using ClassyFire^29^ to annotate each molecule, we found that molecules selected by our model are enriched in benzenoids, ethers, carboxylic acid derivatives, and organoheterocyclic molecules when compared to the molecules measured by the USDA dataset (Figure 4C).

**Figure 4:**
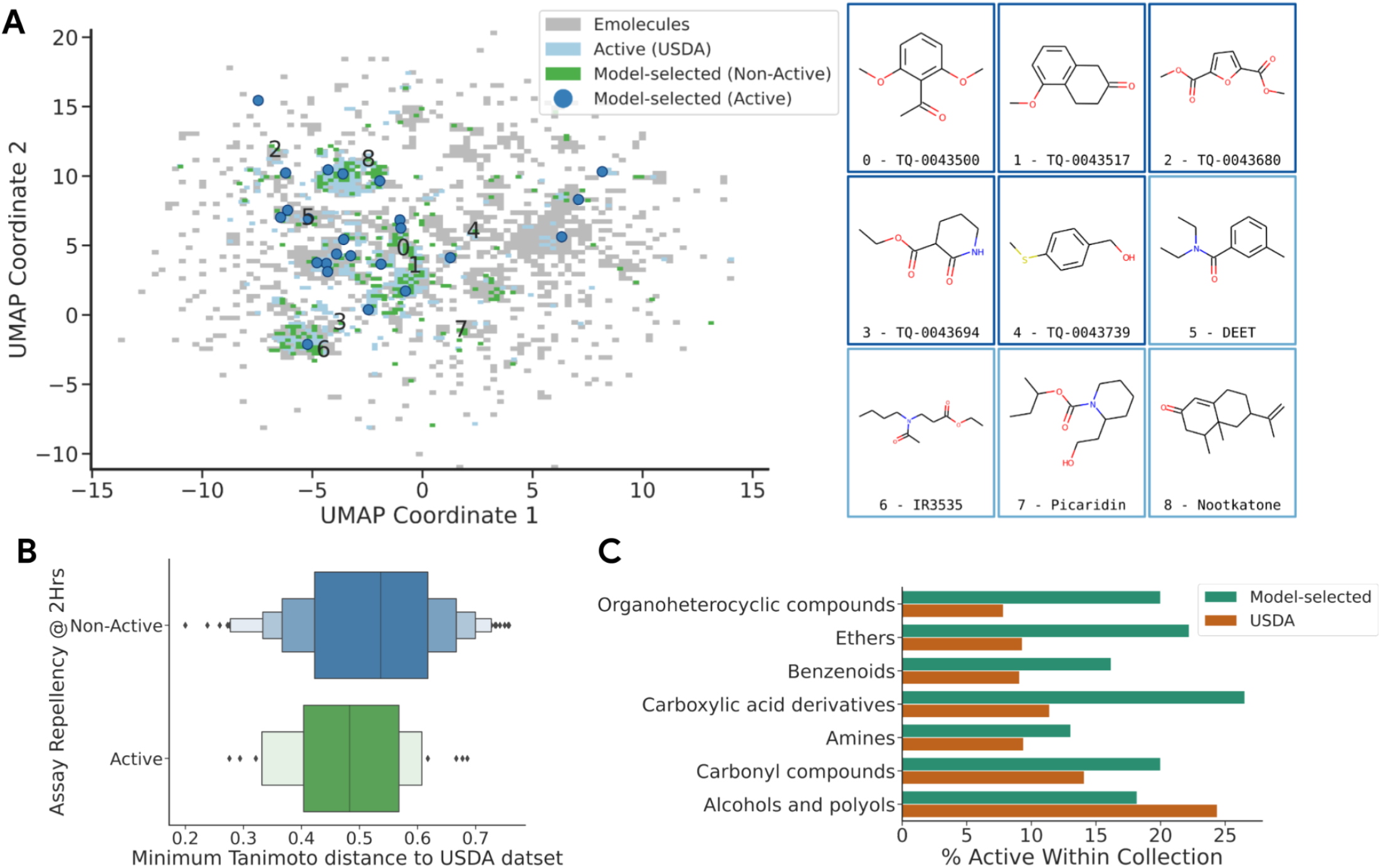
**(A)** The model-selected molecules are distributed throughout the chemical space, with some active molecules found both near and far from USDA clusters. Shown is a UMAP embedding of USDA active molecules (light blue), and model selected molecules (dark blue), aligned with the eMolecules library (grey heatmap), using Morgan fingerprint features (r=4, n=2048). The positions of a few high-repellency, model-selected compounds and several known repellents are shown. **(B)** Tanimoto distance of ML-selected candidates to the USDA dataset; molecules were selected to be at least Tanimoto distance=0.2 away from other USDA molecules, with active candidates having a lower median distance away from the USDA dataset (median=0.48) compared to inactive candidates (median=0.54). **(C)** Distribution of ClassyFire classes (Djoumbou-Feunang et al., 2016) in the USDA dataset and the TropIQ selections. TropIQ selections are enriched for organoheterocyclic compounds, ethers, benzenoids, and carboxylic acid derivatives.

### Top candidates show strong repellency in additional applications

While the membrane feeder assay provides a rapid measurement of repellency effectiveness, for real-world applications it is necessary to consider the effect of odorants released by human skin. To assess repellency of hit molecules in the context of host skin emanations, we tested a representative set of our molecules in arm-by-cage experiments (Fig. 5A). To this end, we selected 31 hit molecules that showed ≥75% repellency at a density of 25 µg/cm^2^ at T=120 minutes at least once in the membrane feeder experiments, and 4 molecules with lower repellency activities. When tested at a density of 13 µg/cm^2^ in the arm-by-cage experiments, 43% of the tested molecules perform very well (≥75% repellency) and 67% of those even outperform DEET (>84% repellency) (Fig. 5B). Overall, we observed high correspondence between repellency as measured in the feeder vs. the arm-in-cage assays (r=0.64), with 83% of hits from the former also reaching the hit threshold in the latter (Fig. 5C).

**Figure 5:**
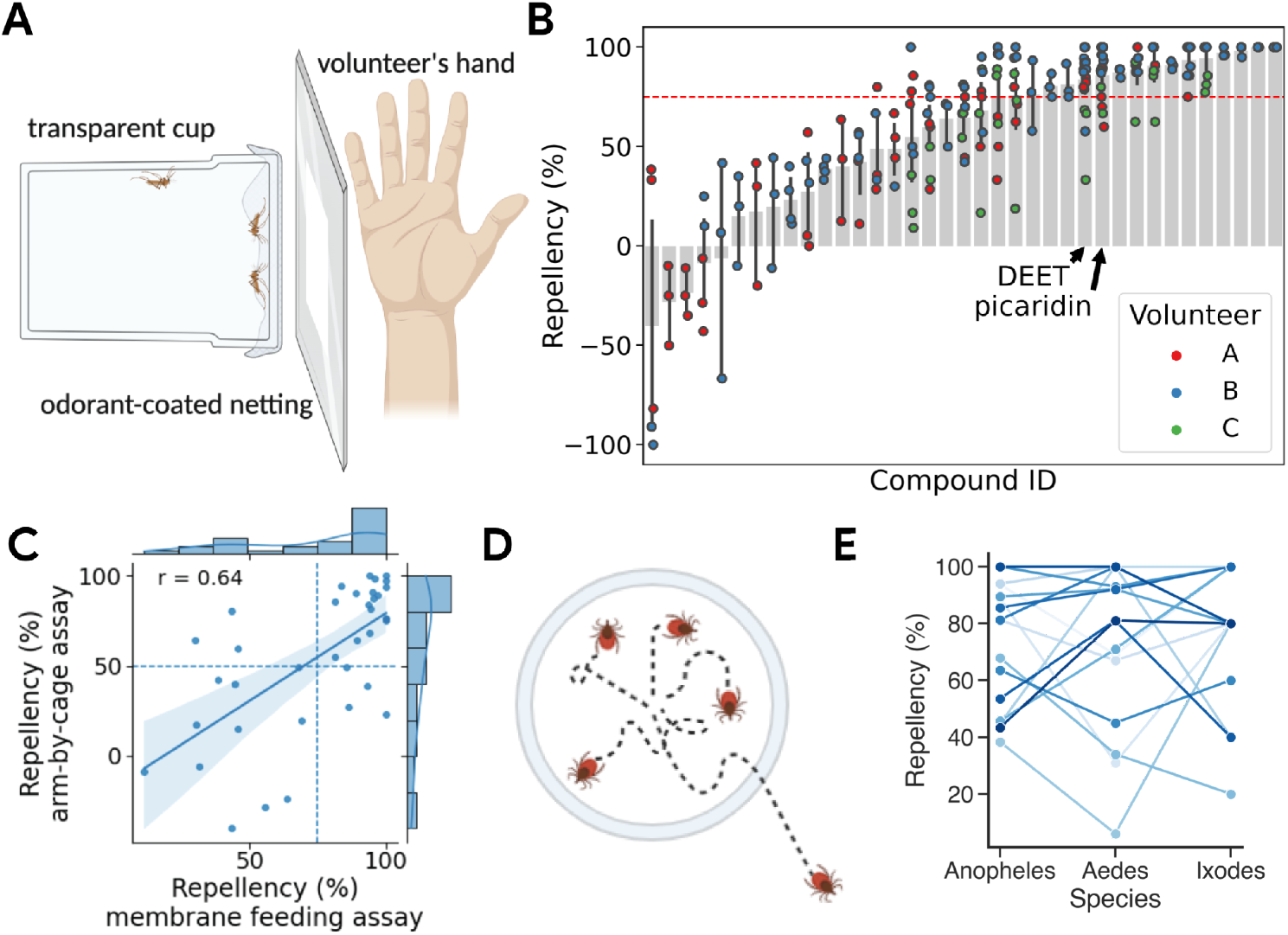
Model-selected and feeder assay validated compounds show high performance across context and species. **(A)** Experimental setup of arm-by-cage experiments on *Anopheles stephensi*. **(B)** Arm-by-cage repellency of molecules previously determined to be repellent in the membrane feeder assay. **(C)** Activities of repellents identified in the membrane feeding assay correlate well with the activity in arm-by-cage assays. **(D)** Experimental setup of *Ixodes scapularis* (tick) repellency assay. Ticks are placed in a repellent-impregnated ring on a heated bed and the number of ticks that cross the ring are counted. **(E)** Repellency of molecules is correlated across species; one line corresponds to one compound.

Our primary assay assessed repellency against *A. stephensi*, but other pest species also carry disease, and there are some known species-specific differences in repellency of known molecules (e.g. IR3535). To address this concern, we selected 16 molecules based on their activity against *A. stephensi*, 9 strong and 7 weak repellents. We then used the original assay to test them against *A. aegypti* and a modified assay (Fig. 5D) to test against *I. scapularis*, the black-legged tick. We observed significant generalization across pest species: 8 of the strong repellents (88%) demonstrated good repellency (>50% repellency) at 25 µg/cm^2^ against *A. aegypti*, and 12 (75%) molecules were active (>75% repellency) at 540 µg/cm^2^ against *I. scapularis* (ED_50_ of DEET ≈120 µg/cm^2^, Fig. 5E).

### Mosquito neural mapping show more patterns to be uncovered

To understand the mosquito neurophysiological perception of the repellents that were selected by our machine learning approach, we conducted two-photon imaging in the primary olfactory center of the mosquito brain, the antennal lobe (AL). The AL comprises dense regions of neuropil, called glomeruli, with each glomerulus encoding a handful of volatile compounds. Experiments using 10 unmated female *A. aegypti* mosquitoes, aged 4-9 days, were conducted. To facilitate repeatable registration of AL glomeruli between preparations and recording of glomerular responses, mosquitoes were immobilized on custom-made holders, and then the upper half of their head capsule was removed to expose the antennal lobes (Fig. 6A). Patterns of glomerular activity were evaluated in GCaMP-expressing mosquitoes (QUAS-GCaMP7s transgene crossed with the brp-QF2 driver line (Zhao et al., 2022)) at three depths in the AL, 15, 30, and 45 µm measured dorsoventrally (figure 6B). A total of 50 regions of interest (ROI), tentatively identified as glomeruli using 3D reconstruction and registration (Shankar and McMeniman, 2021; Wolff et al., 2022), were evaluated for changes in their baseline fluorescence (ΔF/F) after a 2 s exposure to a particular compound (figure 6C). Results of these experiments revealed that certain individual glomeruli showed significant responses to tested compounds, such as DEET, that were locked to the stimulation window before falling to baseline levels after a few seconds (Fig. 6C).

**Figure 6.**
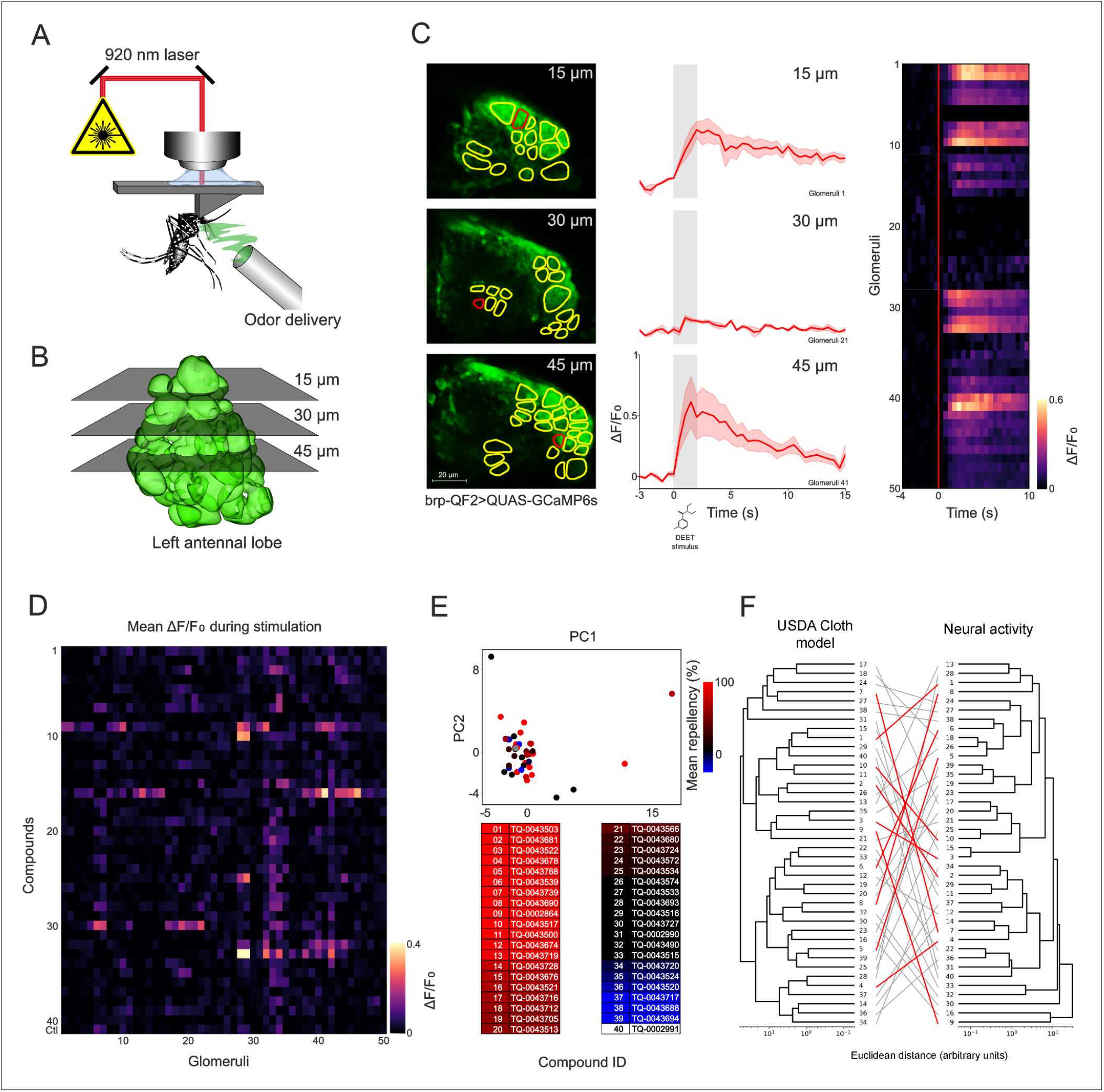
Processing of Olfactory Inputs in the Antennal Lobe of Female *Aedes aegypti* Mosquitoes. **(A)** Female mosquitoes are secured to a plastic holder, where the exposed antennal lobes can be observed from a dorsal perspective, while an olfactory stimulus is released using an odor delivery system. A two-photon microscope is used to observe and record changes in glomerular GFP fluorescence (ΔF/F0) associated with neural responses to odors. **(B)** Frontal view of the left antennal lobe. Neural activity was evaluated across three planes (15, 30, and 45 um dorsoventrally) to maximize glomerular representation. **(C)** (left) Composite calcium images showing the average fluorescence over time of stimulation with DEET at each imaging plane in one individual mosquito. The outlines represent 50 observed glomeruli. (middle) Temporal dynamics of GFP activation in the three sample glomeruli outlined in red when exposed to DEET for 2 seconds (gray region). (right) Glomerular population responses to stimulation with DEET, odor release happens at t=0 (red line) and lasts 2 seconds. **(D)** Mean change in glomerular activity during the two seconds of stimulation across 50 glomeruli and 40 compounds of interest. Compounds are sorted from highest (1) to lowest (39) repellency* as reported from behavioral assays. **(E)** (top) Principal component analysis based on the patterns of neural activity during stimulation. Compounds are color-coded according to their repellency as shown in the table below. **(F)** Hierarchical clustering of compounds tested, based on their predicted repellency according to the USDA cloth model (left) and measured glomerular activity during stimulation (right). Lines were added to evidence the location of each compound in both trees, highlighting in red compounds with over 75% repellency in behavioral assays. *No data available for compound 40 (TQ-0002991)

For each compound, the population of glomerular ROIs showed consistent patterns of neural activation during the time of stimulation (Fig. 6D). From the population-level responses, multivariate analyses were used to examine the relationship between representations of the tested compounds; however, the principal components analysis (PCA) of the mean ΔF/F did not reveal a clear clustering of compounds. To determine if the neural responses correlated with behavior, the Euclidean distances of the neural representations were correlated with the behavioral and cheminformatics data. Results showed that the patterns of neural activity correlated poorly with behavioral data (Spearman’s rho: -0.22; P-value = 0.19), as visualized by color-coding PCA dots based on repellency (Fig. 6E).

Similarly, neural activity in the glomeruli observed exhibited a weak correlation with the neural network representation from the portion of the USDA model trained on the USDA cloth-*A. aegypti* task (Spearman’s rho: 0.29; P-value = 0.0678). This suggests a tentative relationship between the predicted repellency of the compounds tested and their corresponding neural activation patterns. To show the relationships, or lack thereof, between our GNN model, glomerular representations, and repellency, hierarchical clustering analyses were implemented on the chemical fingerprints and glomerular activity patterns, and the behavioral repellency was shown as the lines linking the two dendrograms, demonstrating that strong repellents (red lines) can evoke diverse olfactory responses that are not always correlated to one another (Fig. 6F). Further analysis also showed no correlation between neural activity and chemical fingerprints, as determined by Tanimoto distances to DEET (Spearman’s rho: 0.05; P-value = 0.78). Our results show that tested compounds evoke a diverse set of neural activity patterns, ranging from unique to overlapping. Such diversity of glomerular representations, particularly among strong repellents, suggests that they may be activating both unique and similar olfactory channels in the mosquito. The pattern of channel responses is not fully explained by chemical fingerprints or by our model representations, and would be an important avenue of research for elucidating neural activity.

## Discussion

We developed and validated novel methods for identifying potential repellent molecules for vector control of deadly human and animal diseases. First, we digitized a historic dataset rich with an unprecedented volume of relevant repellency data covering thousands of molecules. Second, we applied and refined a deep learning model architecture to learn the mapping between molecular structure and repellency in this dataset. Third, we used a high-throughput experimental assay to prospectively validate predictions from this model, and to conduct active learning to iteratively improve model predictions. Finally, we showed that these predictions identify new repellent candidates in underexplored regions of chemical space, and that some of these molecules show applicability across real-life context and across pest species. We further assessed neural activity in the antennal (olfactory) lobe of mosquitoes in response to exposure to a subset of these compounds, evaluating whether such activity correlated with our graph neural network models representation. We present this approach as a promising direction to identify next-generation repellents, utilizing historical datasets and machine learning applied to new mosquito assays.

Despite containing a surprisingly large quantity of relevant repellency data, the USDA dataset has remained underused, garnering only ∼200 citations in the last 50 years. This surely stemmed in part from the limited visibility and accessibility of the data during most of this period, where it was accessible only via paper handbooks in physical libraries. The Google Books digitization project scanned these handbooks, making images of the data visible to anyone with an internet connection. However, many of the chemical names contained there-in were archaic or ambiguous, and so could not be effortlessly mapped to chemical structures; the repellency values themselves were also not machine readable. The manual curation and digitization that we performed was the last step to unlock the power of these historical records. The general pattern of connecting diffuse experimental records to support larger modeling efforts and meta-analyses continues to bear fruit^30,31^.

How important were these data? Machine learning is data-driven, and frequently suffers from “cold start” problems; deep learning models are especially data-hungry, and finding enough data to train them to state-of-the-art performance can be a major challenge. The USDA dataset solved this problem by allowing us to train a draft model, which we were then able to build upon using data from a modern experimental assay. Several previous efforts to identify new repellents using machine learning have used only several dozen similar molecules to train their models^15–17,32^. A larger slice of the historical dataset (∼2000 molecules) has been used to train a neural network model to both predict repellency and verify the repellency of known repellents^33^. Recently, larger datasets are becoming available for *receptor-targeted QSAR (RT-QSAR)*^*34,35*^, but until this current work, no machine-readable large-scale datasets have been available for BT-QSAR.

Most previous publications validated their repellency models only retrospectively by predicting the activity of known repellents, rather than prospectively^36^ by using the model to identify new molecules with repellency behavior. This typically leads to overestimation of predictive performance of new repellent candidates. By contrast, we collected assay data for prospective validation of the model, and further used this data in an active learning loop to refine the model, showing continued improvement in predictive performance as new data was collected.

Prospective validation has been used in the past to discover new repellent molecules: Picaridin was discovered at Bayer using pharmacophore modeling^6^, and a small set of acylpiperdines were discovered using neural networks trained on a small subset of USDA data^17^. However, these novel repellents have typically been structural near-neighbors of existing repellents. By contrast, our model-selected candidates cover a much wider range of structural classes than previous repellency discovery attempts, facilitating our discovery of molecules with repellency activity greater than DEET even at 2 hours after application, and a subset that have repellency efficacy when tested in the presence of attractive human skin emanations.

Machine learning, and particularly deep learning, is yielding impressive advances in applications in chemistry. Several academic and industrial groups have used deep learning models to screen for new molecules with desirable properties, such as antibiotic activity or protein binding affinity^34,37–39^. The methods outlined in this paper can also be applied to other disease vectors, other classes of behavior-modifying molecules, and more broadly to enable hit discovery in arbitrary chemical applications. Further approaches could train models to use mosquito neurophysiological response data in addition to behavioral assay data to potentially identify new classes of components. Future work on imposing additional filters or modeling steps to satisfy additional criteria related to safety, biodegradability, odor, and skin-feel, will be required to develop next generations of pest control compounds.

## Methods

### Mosquitoes and ticks

Both *Anopheles stephensi* and *Aedes aegypti* mosquitoes were maintained on a 5% sugar solution in a 26 °C environment with 80% humidity, according to standard rearing procedures. Adult *Ixodes scapularis* ticks were maintained in a 26 °C environment with 90% humidity.

### Mosquito behavioral assays

Before each membrane feeding assay, 10-20 female *Anopheles stephensi* or *Aedes aegypti* mosquitoes (3-5 days old) were transferred to a paper cup covered with mosquito netting. The mosquitoes were denied access to their normal sugar solution 4-6 hours prior to the feeding assay. 30 µl of test molecule, dissolved in ethanol, was pipetted on a piece of mosquito netting (3×3 cm) and allowed to dry. To ensure a regular and standardized airflow over the samples, a gastronorm tray (½ 200mm) equipped with a computer fan (80×80×25mm, 12V, 0.08A) was placed over the samples. After a specified time of evaporation (e.g., 2 hours), the sample was placed on top of the cup containing the mosquitoes. The cups were then placed under a row of glass membrane feeders containing a pre-warmed (37 °C) blood meal. The mosquitoes were allowed to feed for 15 minutes. The number of fed and unfed mosquitoes were then recorded.

For the arm-by-cage assays, 30-50 female *Anopheles stephensi* mosquitoes were transferred to an acrylic cup (150×100mm) covered with mosquito netting. 1 mL of test molecule (0.5% w/v), dissolved in ethanol, was pipetted on a piece of cheesecloth (6×9 cm) and taped to an acrylic panel (6mm thick) with a cutout and allowed to dry. A panel with an untreated piece of cloth was then placed next to the acrylic cups containing the mosquitoes and a volunteer placed his hand against the panel for 5 minutes. The mosquitoes were filmed and the maximum number of mosquitoes landing simultaneously was recorded. This was then repeated with a piece of treated cloth and the number of landings was normalized to the control, which is the ethanol solvent alone. All arm-by-cage assays were designed and run by TropIQ.

### Tick behavioral assays

The setup of the tick repellency assay is shown in figure 5D. The assay consists of a heated (37ºC) aluminum plate (235 × 235 mm) that is painted white. Before the test, 750 µl of test molecule, dissolved in ethanol, is pipetted on a ring of filter paper (OD = 150 mm, ID = 122 mm). The ring is then transferred onto the heated plate and 5 *Ixodes scapularis* ticks are placed in the center. The ticks are monitored for 5 minutes and the number of ticks that cross the filter paper are counted. Repellency is expressed as the percentage of ticks that did not cross the filter paper.

### Calcium Imaging

Odor-evoked responses in the Ae. aegypti antennal lobe (AL) were imaged using transgenic mosquito lines, specifically the offspring of brp-QF2 and QUAS-GCaMP7s, referred to as brp-QF2>QUAS-GCaMP7s progeny (Zhao et al., 2022). We evaluated neural activity in the antennal lobe of 10 unmated female mosquitoes, aged 4-9 days. Each mosquito was anesthetized on ice and then transferred to a Peltier-cooled tethering station kept at 4 °C, where it was secured to a custom stage using ultraviolet glue (Lahondère et al., 2019; Melo et al., 2020). The dorsal half of the head capsule was carefully removed using fine forceps to expose the antennal lobes. Throughout the dissection and imaging process, the brain was continuously superfused with physiological saline at 21 °C (Vinauger et al., 2019).

We tested a panel of 40 compounds of interest, along with two positive controls, known to be highly attractive to female Ae. aegypti (see supplementary information). Test compounds were diluted in ethanol (200 proof; Sigma) to a 1:1000 v/v concentration for positive controls, and to 1:5 v/v concentration for the compounds of interest due to their low volatility. Odor stimuli were presented for 2 seconds and separated by intervals of at least 120 seconds to avoid receptor adaptation. Odor cartridges were replaced every three uses to prevent decrease in stimulus concentration.

Calcium-evoked responses in the AL were imaged using the Prairie Ultima IV two-photon excitation microscope (Prairie Technologies) and Ti-Sapphire laser (Chameleon Ultra; Coherent) set at 1910 mW power. Time series were collected from a 105.6 µm × 79 µm plane at 2 Hz, with a line period of 1 ms. For each odor stimulus, images were acquired for 25 s, starting 10 s before the stimulus onset. To maximize glomerular representation across preparations, experiments were performed at 15, 30, and 45 µm depths from the dorsal surface of the AL. Following each experiment, the AL was scanned dorsoventrally at 0.1 µm intervals to generate a Z-stack that would help provide glomerular assignment and consistent registration across preparations.

We defined, mapped, and registered 50 regions of interest (ROI) as tentative glomeruli, based on odor-evoked responses, location, and morphological features using available AL atlases (Shankar and McMeniman, 2020; Wolff et al., 2023) and Z-stack-based reconstructions in Amira software (v. 6.5, FEI Houston Inc.). Time series generated were analyzed in ImageJ software (version 1.54f, National Institutes of Health, USA; [http://imagej.org]) where ROI were delineated according to glomerular assignments. Calcium responses, indicated by changes in fluorescence (ΔF/F), were time-stamped and synchronized with the odor stimuli. Glomerular responses were aggregated into a neurological representation matrix by extracting from the normalized time series the mean ΔF/F during stimulation for each glomerulus in response to each compound. The resulting dataset represented the glomerular ensemble’s collective response and allowed us to compare neural activity patterns using dimensionality reduction analyses.

### Historical dataset preparation

The scanned versions of the USDA datasets, available from Google Books, were converted into a machine-readable format. Chemical structures (Simplified Molecular-Input Line-Entry System, or SMILES) ^40^ were assigned to each single molecule entry in the dataset. The raw PDFs of the two repellency handbooks^41,42^ used to create the USDA dataset are available on Google Books. For this study, the PDFs were converted to png files, then sliced by rows according to bounding boxes drawn by curators. The row sliced images and the full page images were provided to a third-party curation service, who transcribed the chemical names as SMILES and corresponding assay results. Post-processing analysis and evaluation of a random sample of 150 entries suggest an error rate of <5% in the chemical structures. The final dataset resulted in 18,886 data points on 14,187 molecules. This includes the results on two assay setups, one testing the effectiveness of the candidates on cloth, the other on human skin, and also two different mosquito species (*Aedes aegypti* and *Anopheles quadrimaculatus*); all four combinations of these two species and conditions were used in this study. USDA dataset labels in the source material were repellency ratings given as integers from 1 (worst) to 5 (best).^41^

### USDA Dataset Modeling and Representation Learning

Each of the USDA tasks was split into a 70:15:15 train/validation/test split such that molecules were assigned to the same split across all tasks; in particular, if a molecule is in the training set for one task, it was also in the training split for the other tasks for which there was a measurement. Molecules in the USDA dataset that were also used in the pre-USDA phase (Batches 1-3, see Supplementary Batch notes) were excluded from the USDA training sets. Iterative stratification over the label classes across each task was applied to balance the labels in the training/validation/test splits for each task.

Graph neural network models (GNNs) were trained on each of the four mosquito repellent tasks from the USDA dataset. Each model provided predicted probabilities of the class label and combination class labels; specifically, the model predicted the probability of the class label being: [1], [2], [3], [4], [5], [1 OR 3 OR 4 OR 5], [3 OR 4 OR 5], [1 OR 4 OR 5]. AUROC performance on the [3 OR 4 OR 5] label objective was used to optimize the models. The graph neural network used message passing layers (MPNN^44^), with a max atom size of 45, 30 atom features, and 6 bond features. Hyperparameter selections were made using the Vizier^43^ default Bayesian optimization algorithm over 300 trials.

The USDA learned representation was constructed from the outputs of the frozen ensemble model of the best 50 models from hyperparameters trained on the USDA dataset. For the last batch of selections, the models used to create the ensemble model ranged in AUROC performance from 0.872 to 0.881.

### Model Training on Membrane Feeding Assay Data

To train the models for activity in membrane feeding assays, assay results were binarized: a positive label for repellency activity was defined as >90% at T=2min at 25 µg/cm^2^, and >75% for T=120min. For model evaluation and hyperparameter selection, the dataset was split into a 70:30 train/test split, using iterative stratification to balance the label classes. The model trained on the USDA dataset was used to generate specialized representations for the molecules. A two-layer neural network model with 256 nodes was used to predict the binarized activity label given the molecule; the hyperparameters of this model were selected with grid search. At inference time, to make predictions on new candidates, the model was retrained using the entire dataset.

### Molecule Selection

We began by filtering molecules listed in the eMolecules catalog -- which contains ∼1 million commercially available molecules -- for atom composition (C/N/O/S/H only), price (<$1000 per 10 grams), purity (>95%), and availability (<4 weeks lead time). We utilized a toxicity filter to remove potentially harmful molecules, according to a toxicologist-recommended protocol. In this protocol, we classified molecules by their mutagen / Cramer class using ToxTree, calculated their vapor pressure at room temperature, and then compared the likely exposure air volume to OSHA daily exposure limits for the corresponding toxicity class. We removed likely odorless molecules according to water-soluble (cLogP < 0) and nonvolatile (boiling point > 300 C) criteria. We manually removed molecules that were likely to degrade or react under our experimental conditions. After training the assay model, molecules were selected such that they had a prediction score above an f1 optimized cutoff score, and then selected such that they had a Tanimoto similarity of <0.8 from other selected molecules and the USDA dataset. A minimum solubility threshold of 10 mg/ml in absolute ethanol was used as a last criterion. Molecules with an ethanol solubility below the threshold were abandoned. Detailed selection criteria for batches are reported in the Supplemental section.

## Author Contributions

JNW, DMA, KMG, ABW curated and digitized the USDA dataset; JNW, BKL performed data cleaning and spotchecking of the dataset. MV and KJD designed the mosquito assay and tick assay experiments; MV, LB, MWV, and RWMH performed the mosquito assay experiments; MV and MWV performed tick assay experiments. CR, JAR, MV, and JNW designed the mosquito calcium imaging repellent panel; CR performed the experiments and data analysis. JNW designed the models with assistance from BS-L, BKL, and WWQ. JNW, MV, BS-L, RCG performed data analysis. JNW, MV, RCG wrote the manuscript. ABW and KJD conceived the project. All authors contributed to editing the manuscript.

## Acknowledgements

We wish to thank Laura Pelsen-Posthumus for technical assistance in mosquito rearing, and Geert-Jan van Gemert and Pascal Miesen for provision of mosquitoes. We thank Hans Dautel for help with the design of the tick repellency assay. The authors thank Lucy Colwell, James Thompson, Max Bileschi, and David Belanger for critical reading of the manuscript. We also thank Sameer Kulkarni for assistance with OCR processing and Jonathan Brecher for his insights on transcribing structures from historical chemical names. We thank Ben Adlam, Jasper Snoek, and the Cambridge Brain Research team for giving helpful feedback during our discussions. Some of the figures were created with BioRender.com. We also thank the Bill and Melinda Gates Foundation for their generous support of this work, and the support from the National Institutes of Health under award R01AI148300 (JAR), which helped fund some of the calcium imaging work.

## Data Availability Statement

With this publication, we make available in csv format the repellency results and the calcium imaging results for the 40 compounds tested with the calcium imaging assays. The digitized version USDA dataset^4,5^, code used to train the mosquito repellency models, and the models, are the intellectual property of Osmo, and are not available with this publication. The full repellency results for all 317 selected repellents are the intellectual property of TropIQ, and are not available with this publication.

## Current Author Affiliations

## Supplemental Information

### Batch Selection Notes

#### Pre-USDA batches

Batch 1: 29 molecules were selected containing known repellents and controls based partly on the literature of Oliferenko et al^45^, Carey et al^46^, and Xu et al^47^.

Batch 2: The Principal Odor Map reported in Qian et al^26^ was used to embed molecules and a k-nearest neighbors model (k=10) was trained on activity for 31 molecules on the membrane feeder assay at T=2min. 39 molecules were ordered and tested.

Batch 3: 23 molecules found in USDA dataset^5^ were tested on the membrane feeder assay. The molecules were distributed across mosquito repellency classes assigned in the USDA dataset.

#### Post-USDA batches

Batch 4: *Assay model:* A Graph neural network (GNN) model was trained on the Skin Repellent assay results on *Aedes aegypti* from the USDA dataset^5^, containing 6,111 data points, using the setup described in the Methods section USDA Dataset Modeling. This model was used to run inference on the eMolecules library. An initial selection of 80 molecules such that the molecules had a predicted score of higher than 0.4 and a Tanimoto similarity of 0.8 or less from other selected molecules. These molecules were further filtered for toxicity and solubility, resulting in a final set of 33 molecules.

Batch 5: *USDA learned representation*: A graph neural network model was trained on the USDA mosquito repellency data found in King, 1954.^41^, using the results from all four experiments (Cloth - *A. aegypti*, Cloth - *A. quadrimaculatus*, Skin - *A. aegypti*, Skin - *A. quadrimaculatus*), for a total of 18,866 datapoints. The USDALR are the outputs of the fixed ensemble model constructed by averaging the predictions by the four models trained on the USDA tasks. Assay model: A random forest model was trained to use the inference from the ensemble model to predict binarized membrane assay repellency activity. The membrane assay dataset used to train this model had repellency time activity at T=2min on 142 molecules. This assay model was run on all of eMolecules; 33 molecules were selected such that they had predicted scores in the top 80% and had a Tanimoto similarity of not greater than 0.8 from other previously tested molecules and the other selected candidates.

Batch 6: *USDA learned representation*: A graph neural network model was trained on all four USDA mosquito repellency tasks found in King, 1954^41^, the same as used in Batch 5. The USDALR are the fixed outputs of the ensemble model constructed by concatenating the predictions from the four USDA models. *Assay model*: Two neural network layers of size 256 were appended to the USDA model and trained activity of 320 molecules on the membrane feeder assay dataset for T=2min. When training on the assay dataset, the USDA model weights were frozen; that is, only the neural network layers were tuned. *Molecule Selection*: Molecule selection was performed in two waves. In the first wave, 89 molecules were selected such that they had a predicted score above the 55th percentile and such that they had a Tanimoto similarity of 0.8 or less from USDA molecules, previously tested molecules, and other selections. In the second wave, 75 molecules were selected, with the prediction threshold lowered to the 40th percentile and the same structural similarity filters.

Batch 7: *USDA learned representation*: The outputs of a fixed ensemble model of the best 50 graph neural network models trained on the yellow fever cloth task of the USDA dataset. The models in the ensemble ranged in AUROC performance from 0.872 to 0.881. Assay model: USDA model combined with two neural network layers of size 256 nodes. This model was trained on the activity of 402 molecules on the membrane feeder assay at T=120min. 72 molecules were selected with a prediction score in the 75th percentile, such that they have a Tanimoto similarity of 0.8 or less from previously tested molecules, USDA molecules, and other selected molecules.

